# From foodwebs to gene regulatory networks (GRNs) - weak repressions by microRNAs confer system stability

**DOI:** 10.1101/176701

**Authors:** Yuxin Chen, Yang Shen, Stefano Allesina, Chung-I Wu

## Abstract

More than 30% of mRNAs are repressed by microRNAs (miRNAs) but most repressions are too weak to have a phenotypic consequence. The diffuse actions have been a central conundrum in understanding the functions of miRNAs. By applying the May-Wigner theory used in foodweb studies, we show that i) weak repressions cumulatively enhance the stability of gene regulatory network (GRN), and ii) broad and weak repressions confer greater stability than a few strong ones. Transcriptome data show that yeast cells, which do not have miRNAs, use strong and non-specific mRNA degradation to stabilize their GRN; in contrast, human cells use miRNAs to increase degradation more modestly and selectively. Simulations indicate that miRNA repressions should be distributed broadly to >25% of mRNAs, in agreement with observations. As predicted, extremely highly expressed genes are avoided and transcription factors are preferred by miRNAs. In conclusion, the diffuse repression by miRNAs is likely a system-level strategy for enhancing GRN stability. This stability control may be the mechanistic basis of “canalization” (i.e., developmental homeostasis within each species), sometimes hypothesized to be a main function of miRNAs.

## Introduction

Large networks are inherently unstable. Gene Regulatory Networks (GRNs) are large and may thus potentially be unstable, particularly with the low abundance of stabilizing features such as immediate negative-feedback loops (e.g., predator-prey interactions). In this study, we address the stability of large RNA networks by applying the May-Wigner theory, (1-3) which has been extensively used on species interaction networks. Network stability is defined here as the speed of restoring transcript abundance across the transcriptome when it is perturbed. The advantages in using this theory on GRNs are many. For example, the size, composition, connectivity, interactive strength and decay rate in GRNs can all be empirically measured (4-8). In this study, we compare GRNs of human and yeast, with and without microRNAs, respectively, for their stability control.

MicroRNAs (miRNAs) are a class of small regulatory RNAs, which degrade mRNAs and repress translation. For regulatory genes, miRNAs seem paradoxical for two reasons: i) exclusive down-regulation of their direct targets; ii) broad and weak repressions of hundreds of target genes. In comparison, transcription factors (TFs) up-and down-regulate their targets with comparable frequencies (6-8) and often exert strong effects on gene expression as part of a larger transcription complex (9-12).

There are two contrasting views on miRNAs. In the conventional “phenotypic” view, miRNAs repress targets to effect phenotypic changes. Since the vast majority of the repressions are too weak to exert a measurable influence(13, 14), only a small number of targets are considered functional in this view. The rest is assumed to be noises(13, 14). The issue is contentious even for the same miRNA and phenotype(15, 16). An accompanying study (Liufu et al.), which analyzes multiple miRNA targets and phenotypes concurrently, also reports miRNAs’ control of phenotypes to be redundant and incoherent.

In an alternative view, miRNAs function to stabilize gene regulatory circuitry (17-21) and, indirectly, maintain phenotypic homeostasis. Therefore, in one view, miRNAs drive phenotypic changes but, in the other view, keep phenotypes stable (17-21).

Since Waddington first proposed that living organisms must develop along defined paths under irregular input signals, the molecular basis of “canalized development” has attracted much interest (19, 20, 22-24). Heat shock proteins and miRNAs have both been suggested to play that role, at the level of protein and GRNs, respectively(25-27). In the canalization view, miRNAs are often part of regulatory circuits that contribute to stability(28-31). As small circuitries are embedded in larger GRNs, approaches have been developed to model large networks of RNA:RNA cross-talks, in which miRNAs play a central role(32, 33). Such an approach is expanded in this study.

### I. Diffuse actions of miRNAs

We first present mRNA repressions by miRNAs in human cells. Unlike previous analyses(34-42), this study pays special attention to weak interactions, which will later be subjected to mathematical interpretations.

#### Number of targets

We examine 178 conserved miRNAs in human cells for their target sites following the common protocol (Fig. 1A; see Methods). Random seeds with the same CG content serve as the control. If all potential targets are counted, the median number of target genes would be 694, more than 60% higher than the control. The numbers for the moderately and highly conserved targets are 473 and 114 respectively 64% and 185% higher than the control While highly conserved target sites are generally considered more reliable, Xu et al. (2013) have shown that weakly conserved targets are also evolutionarily significant. Hence, the number of targets per human miRNA is likely to be between 100 and 500 (43, 44)(Fig. S1-A,B).

**Figure 1.**
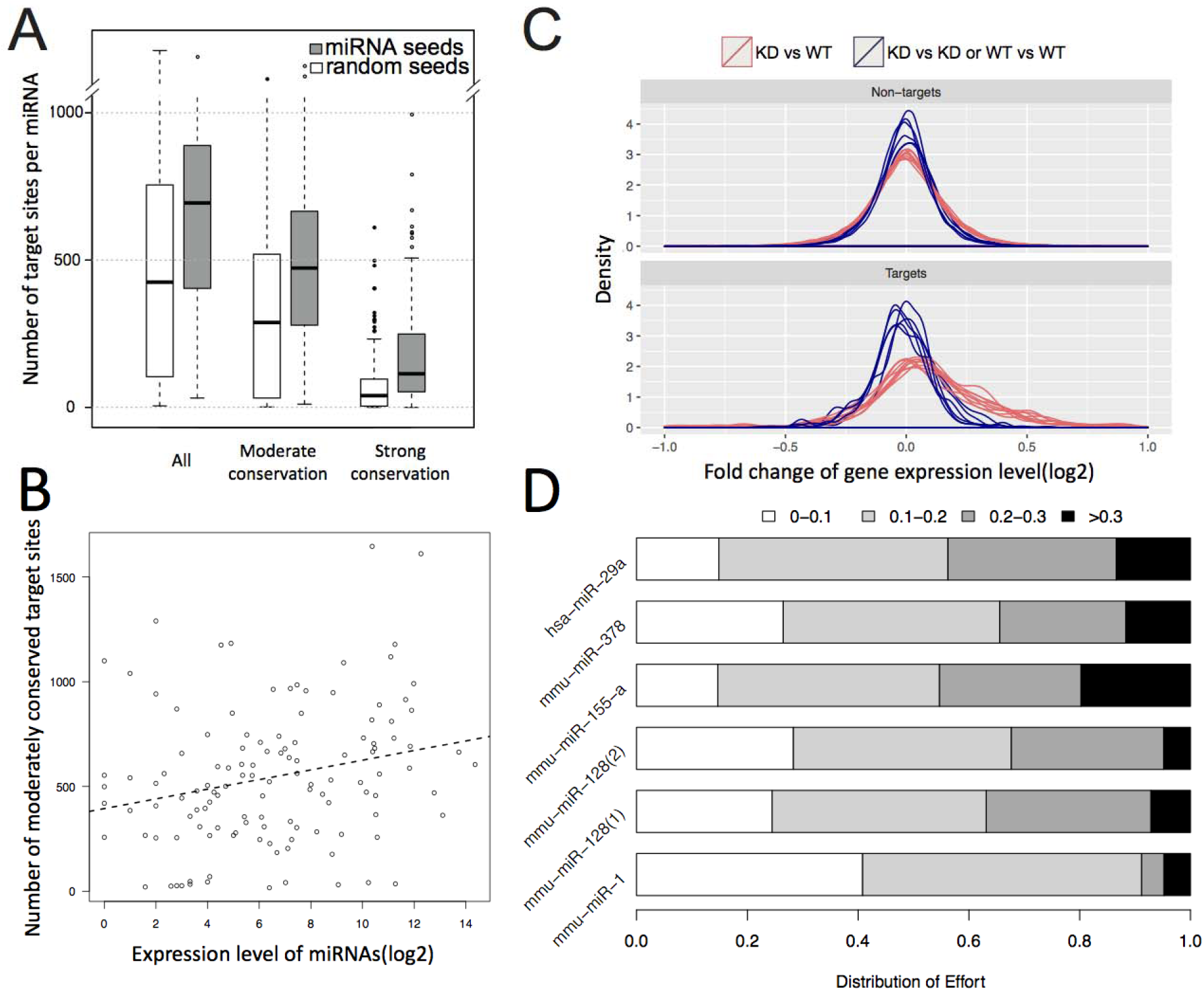
Predicted target number in relation to the observed de-repression by miRNA knockout. (A) Number of miRNA target genes predicted by Targetscan (grey bars) vs. control (white bar) based on the shuffled seeds of the same miRNAs. The comparison is done at three levels of evolutionary conservation. (B) Correlation between the expression level of 109 miRNA seeds and the predicted number of moderately conserved targets. The correlation is positive but the slope is very small (see text). (C) Distribution of fold change in the expression of target genes in miRNA (hsp-29a) knockout lines between experiments and controls (red lines) and between controls (blue lines). The median increase upon miRNA knockout is < 10%. (D) Distribution of effort (DOE) on target repression by each of 6 miRNAs. These efforts are categorized into 4 levels depending on the effect of repression, ranging from <10% to >30%. DOE sums up the repressions across all target genes, weighted by their expression level. Strong repression of >30% generally takes up ∼10% of a miRNA’s repression capacity.

The large number of target sites is even more puzzling for lowly expressed miRNAs. Given their limited repression capacity, these miRNAs might be expected to have far fewer targets. Fig 1B shows the prediction to be qualitatively true. However, the slope of the regression is extremely mild with a decrease of 1/3 in target number when the expression decreases by > 1000 fold(Fig. S1-C,D). Hence, if only strong repressions are functional, then more than half of miRNAs expressed in any tissue would be non-functional.

#### Strength of repression

With hundreds of targets, each miRNA is expected to exert weak effects on most targets. A typical example is given in Fig. 1C which is based on 6 transcriptome datasets from the knockout line of hsa-29a miRNA(Fig. S2). The fold changes of target genes are symmetrically distributed around a peak that corresponds to ∼3% repression. Note that the peak is not at 0%, as is the case for non-targets. Even though hsa-29a is moderately to highly expressed, the degradation of its targets is no more than 5%, on average.

Weak repression could still be noises as long as the weak targeting collectively does not take up much of miRNAs’ total capacity. We therefore measure the fraction of each miRNA’s capacity that is used in weak repression. The distribution of effort (DOE) sums up all repressions of a certain strength, weighted by the expression level of the target gene. Fig. 1D shows that miRNAs use most of their repression capacity to exert small influences on a large number of target genes. Indeed, only ∼10% of the total repression capacity is used for the stronger repression (black bar, Fig. 1D) and the remaining 90% of the effort remains unclear. If we consider miRNAs that are themselves lowly expressed, DOE across all miRNAs would be even more biased toward weak repressions. We next analyze weak repressions in the context of the gene regulatory network (GRN).

### II. GRN stability in relation to expression repression

May (1973) pointed out that large interacting systems are difficult to stabilize, contradicting the belief that large systems are inherently stable. The theory may be particularly suited to GRNs because cell functions depend on transcriptome stability (45) and losing even a small number of genes can have severe consequences (46, 47). While GRNs are periodically perturbed by cell divisions, they lack many stabilizing features like immediate negative feedbacks common in species interaction networks (SINs, e.g., the predator-prey interactions)(1). GRN may thus be considered inherently unstable even though the stability is vital to the cell. Curiously, network stability has been central to SIN studies but has been neglected in GRN analyses.

In a GRN with *N* genes, let *x*_*i*_*(t)* denote the mRNA concentration of gene *i* at time *t*. When the system is at an equilibrium, 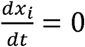 for all *i*’s. Here, we approximate local perturbation near the equilibrium by a linear system (although the system could be non-linear globally):

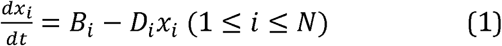

where
*B*_*i*_ = *b*_*i*_ + *S*_*i*_with b_i_ being the basal transcription rate and

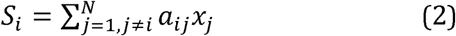

*S_i_* is the aggregate effects of other genes on gene *i* with *a_ij_* being the regulation strength of gene *j* on gene *i*. *D*_*i*_ is the decay rate of the mRNA of gene *i*.

Following the approach of May (1971) and Allesina et al. (2012) for studying SIN stability, we designate the interactions among genes by a matrix, *M*. The diagonal element, *M*_*ii*_, represents the effect of *x*_i_ on itself and the off-diagonal element, *M*_*ij*_, is the regulation strength of gene *j* on gene *i*. M is the Jacobian matrix:

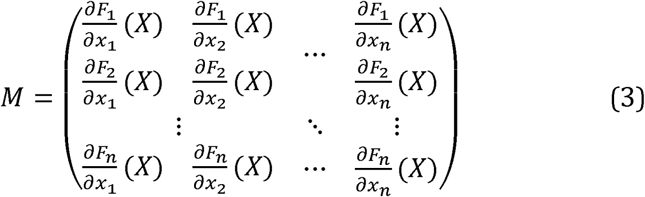

where

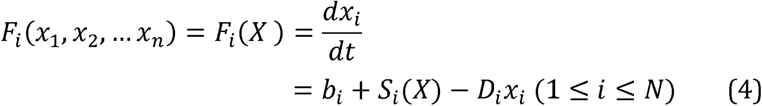

Given Eqs (3) and (4), the elements of the matrix are elements of the matrix are

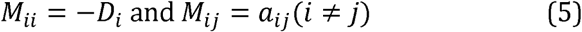

We first consider a network with only one gene (*N* = 1) where the stability condition is

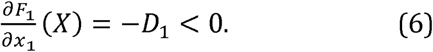

In other words, the slope of *F*_1_ at the equilibrium is negative. In this system of *N* = 1, the local equilibrium is

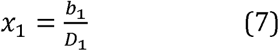

By increasing *D*_*1*_ and *b*_*1*_ in proportion, this system could gain stability without changing the equilibrium and, indeed, the transcription and degradation have been shown to co-evolve (48, 49). Note that the solution of Eq. (7) is a local, rather than global, equilibrium. The latter can be found by empirical means such as RNAseq and the theory is concerned with the local stability near a given equilibrium.

When *N* > 1, the stability of the system is measured in *N* orthogonal directions. The equivalent of *N* negative slopes pertaining to the stability is expressed as *N* negative eigenvalues, which is satisfied if and only if the leading eigenvalue of the matrix *M* is negative. The leading eigenvalue can be approximated as *R* – *D* (Allesina 2012)(1, 50). *R*, a function of the interaction strength (i.e., the off-dianonal elements), is the leading eigenvalue of the matrix *M*_*0*_, which has the same off-diagonal elements as *M* but all diagonal elment elements are 0. 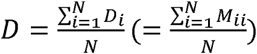 is the average degradation rate. Therefore, the stability condition is

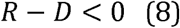

While R and D are usually obtained numerically, an analytical approximation can be derived from Eq. 1 of Tang (2014) when applied to actual transcription data of yeast and mammals (see later sections). Let the connectivity r be the proportion of non-zero *M*_*ij*_’s and let *u* and ***σ*^2^** be the mean and variance of the non-zero off-diagonal elements. When or < 0, and

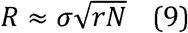

Therefore, the stability condition is approximated by:

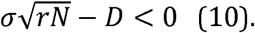

Eq. (10) is suggestive of the roles of miRNAs, which increase *D* by catalytically degrading mRNAs. The degradation can be expressed in two parts

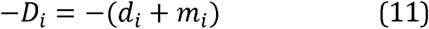

where is the basal decay constant and *m*_*i*_. is the total effect of all miRNAs on the decay of gene Clearly, larger *D*_*i*_’s would make the system more stable.

### III. Comparative anatomy of GRNs with and without miRNAs

Since miRNAs could contribute to GRN stability by increasing *D* (see Eq. 10), we compare human and yeast GRNs by first estimating the diagonal and off-diagonal elements of the interaction matrix, *M*. Yeast cells do not have miRNAs.

#### a. Degradation (the diagonal elements)

The degradation rates of transcripts in many GRNs have been measured, usually by turning off the transcription and monitoring the decay of mRNAs (51, 52). It has been known that the mean half-life for yeast mRNAs is ∼ 15 minutes (4) whereas it is 4 – 8 hours for human mRNAs (5). Fig. 2A and 2B show that the median decay constant (measured in molecules/hour) for yeast mRNAs is 17.4 times larger than that for human mRNAs. Interestingly, when calibrated against their respective cell doubling times of 1.5 and 24 hours, the degradation rates are roughly equal (6.87 vs. 6.33) for yeast and human transcription factors (TFs, red shades of Fig. 2A and2B).

**Figure 2.**
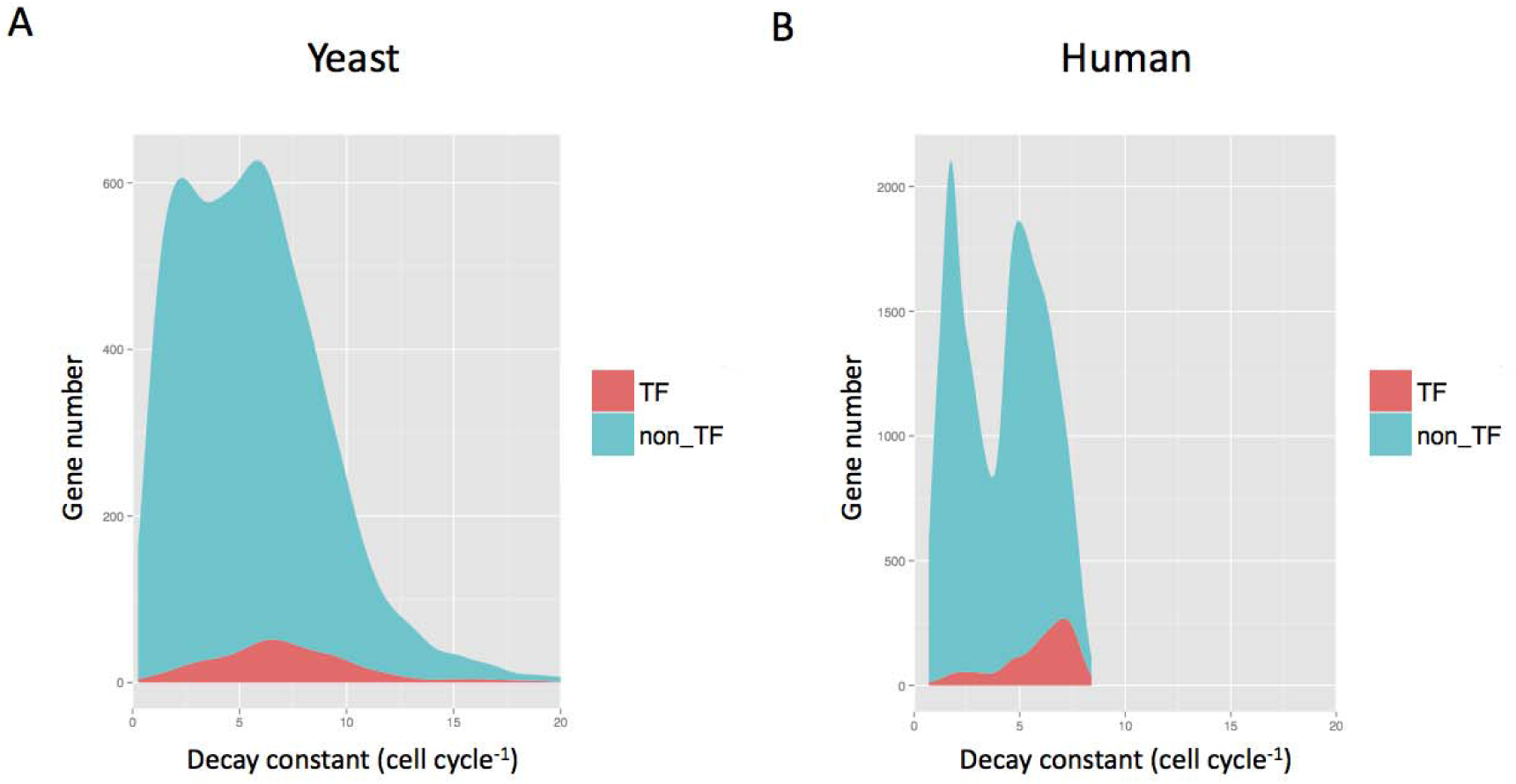
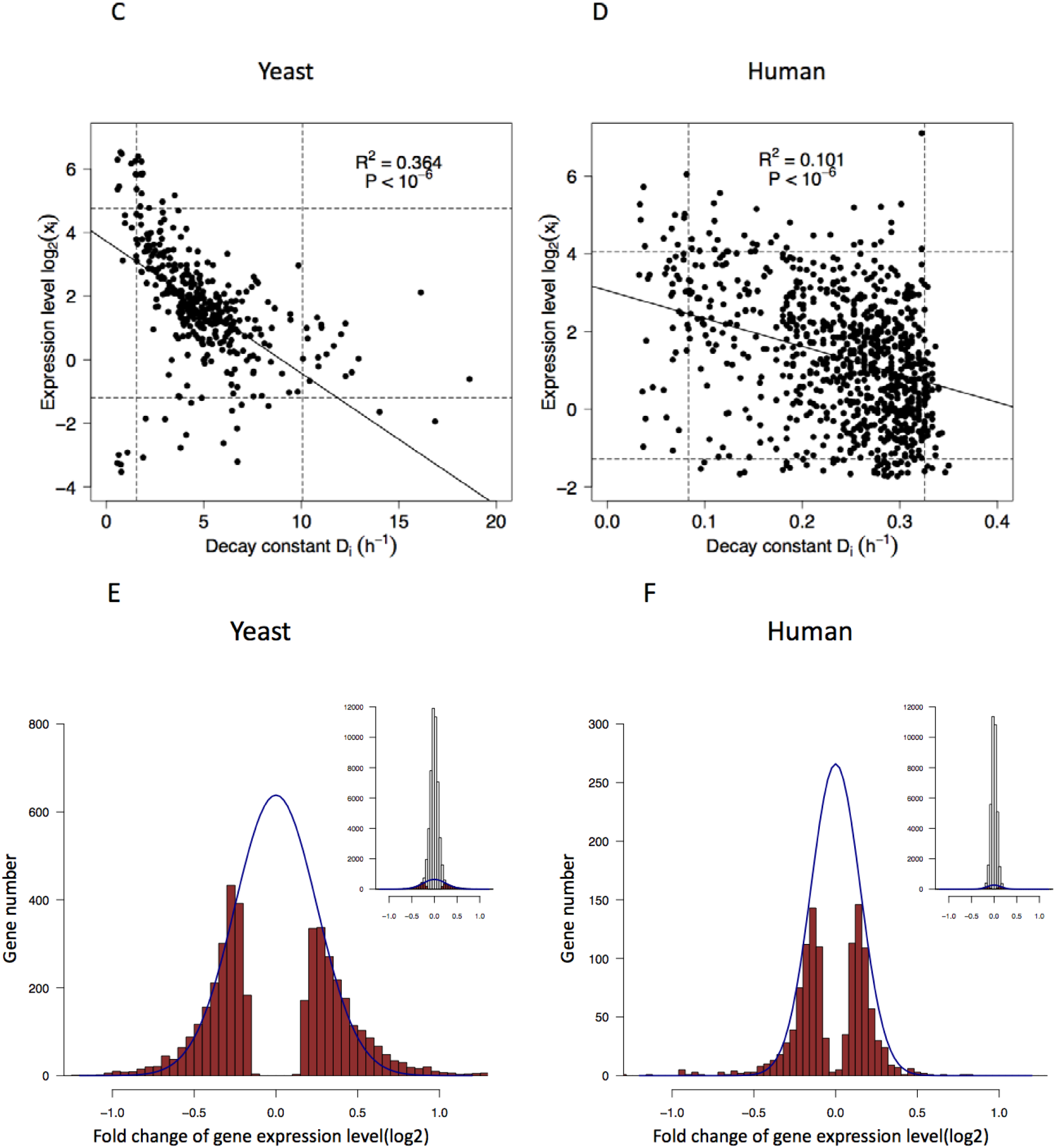
Measurements of degradation and interaction in GRNs. (A, B) Density plot of *D*_*i*_ (mRNA decay constant). TFs (shaded in red) have higher degradation rate than the rest of the transcriptome (blue). The rates shown are calibrated by the respective cell cycle time, by which the two systems are comparable in degradation. In actual time, yeast mRNAs are degraded 17 times faster than human mRNAs. (C, D) The relationship between the expression level of mRNAs and the decay constant. The Y axis spans 3 orders of magnitude while X spans only one order. Hence, the expression level of genes is only marginally affected by the degradation constant. The dotted lines mark 5% and 95% of the distribution. In this restricted range, Y also varies more than X by 10 fold. (E-F) Distribution of the interaction strength between genes in yeast and human GRN. The strength is the change in the abundance of mRNA of gene *i* upon the knockout (yeast) or knockdown (human) of gene *j*. Significant changes (p < 0.001) are marked in red and approximated by a normal distribution marked by the blue line. The inset displays this portion of significantly changed genes relative to all genes.

One might expect the variation in degradation to be driven by natural selection to fine-tune the expression level, *x*_*i*_. The regression of *x*_*i*_.over *D*_*i*_. is indeed significantly negative for yeast and human genes but the correlation coefficient is small (*R*^2^ = 0.364 and 0.101 for yeast and human respectively; Fig. 2C – 2D). Importantly, this is not a simple inverse relationship as *x*_*i*_. spans 3 orders of magnitude and *D*_*i*_. varies by only one order (Fig. 2C-D). Even when we exclude the tails of the distributions (5% on either end), *x*_*i*_.still varies 10 fold more than *D*_*i*_.

The variation in expression, therefore, is largely due to the variation in synthesis rather than degradation (see Supplement on *D*_*i*_ variation). If the many cellular components, including miRNAs and RNA-binding proteins, that function in mRNA degradation do not set the level of gene expression, the question is then “what roles may mRNA degradation play in the GRN”.

#### b. Strength of gene interaction (the off-diagonal elements)

Fig. 1 shows that *D* is much larger in the yeast GRN than in human. Since the smaller *R-D* is, the greater the stability becomes (Eq. 8), yeast GRN could be much more stable than human GRN. Alternatively, yeast GRN might have a correpondingly larger *R* (Eq. 9) and the two GRNs would be comparably stable. We hence analyze the measurements of *M*_*ij*_’s based on experiments that delete or suppress the expression of one transcription factor at a time(53). The TF sub-network is most responsible for the stability of the entire GRN, given its higher position within the hierarchy (see below). The effects of TF deletion/suppression are assayed by transcript analysis (see Methods for details).

We now describe the construction of the yeast GRN. In order to determine the proper size of the network, we rank genes by their expression in the descending order. The set of the most highly expressed TFs with N=356 collectively account for 99% of total mRNAs. N= 356 is the size of the yeast GRN. The procedures for estimating the regulation strength have been widely reported. Several are used here (7, 8) (see Methods).

The distribution of the estimated regulation strength is given in Fig. 2E and its inset. Among all interactions, 4234 are significant with P < 0.001, yielding a connectivity of *r* = 0.076. Fig. 2E shows the significant regulations by the red bars which is approximated by a normal distribution. The non-significant regulations are set to 0. The normal distribution containing all significant interactions is shown relative to the entire set in the inset. In summary, positive:negative regulation is evenly split with a 0.504:0.496 ratio. The mean (*u*) and standard deviation (***σ***) are 0.0144 and 0.432 (see legends). The mean and standard deviation of the absolute value of the interaction strength (|*M*_*ij*_|) are, respectively, 0.379 and 0.207. We note that, in order to construct the GRN, we estimate the mean, variance and distribution of *M*_*ij*_’s. The identities of specific nodes that are connected are not crucial in determining the identify. This aspect of GRN in relation to miRNA function will be discussed.

In the human GRN, the corresponding *M*_*ij*_ distribution is shown in Fig. 2F where *N* = 746 and *r* = 0.031. The positive : negative split is 0.53:0.47, *u* = −0.0322 and ***σ*** = 0.244. The mean and standard deviation of |*M*_ij_| are 0.195 and 0.151 (see Methods). A presentation of a small portion (50x50) of each of the two GRNs is shown in Fig. 3A. The comparison visually portrays the difference in connectivity between human and yeast GRNs (*r* = 0.031 vs. *r* = 0.076) with the latter being denser. Therefore, while human GRN is larger than yeast’s (*N* = 746 vs. *N* = 356), the number of connections per node, —, is very similar with *Nr* = 23.1 vs. 27.1. The effective sizes (54) are hence similar between the two GRNs. The interaction strength in the human network appears weaker, but only mildly so.

**Figure 3.**
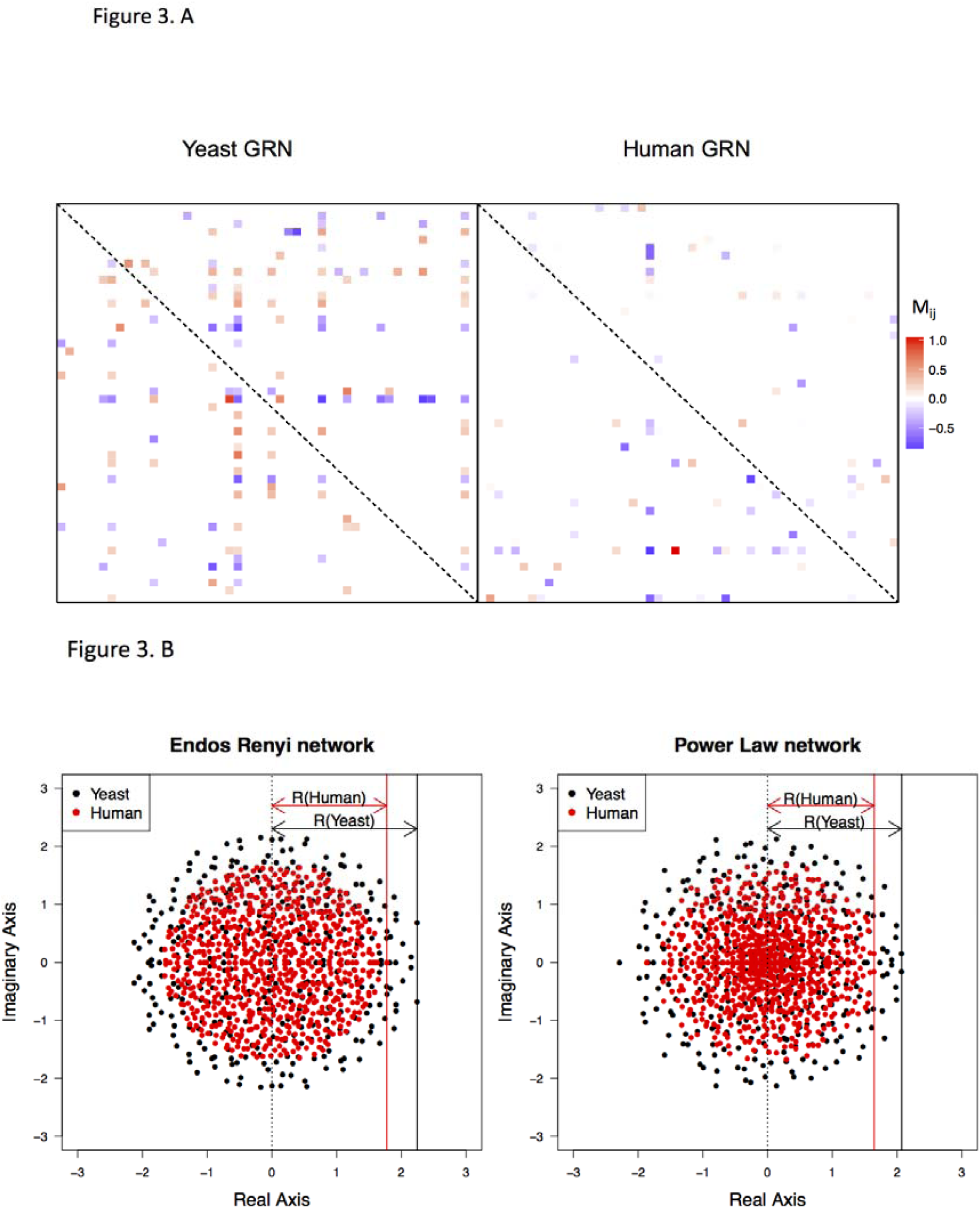
The interaction matrix (*M*) and its eigenvalues. (A) A random 50x50 block of the interaction matrix, *M*, is shown for yeast and human GRN. Off-diagonal elements that are significantly positive or negative (see Fig. 2E-F) are indicated by a color of the heatmap. Diagonal elements are not shown (marked the dashed line). Note that the yeast GRN is slightly denser than human GRN. (B) Distribution of the eigenvalues of GRNs, which are complex numbers with a real and imaginary part. The GRN is constructed with the parameters obtained from the measurements of Fig. 2 with the off-diagonal elements follow a normal distribution (Fig. 2E-2F) and the diagonals set to zero. Two different network structures (random network and power law) are modeled but the outputs are similar. The leading eigenvalue, marked by a solid vertical line, has a value of *R* (Eq. 8 and Eq. 10). The GRN stability requires that *R-D* < 0.

In both networks, significant regulations are not randomly distributed among nodes as a small fraction of nodes are disproportionately more connected than the rest (55, 56). Fig. S3 presents the distribution of in-degree connections, or the number of significant regulations going toward a node as required in Eq. (2). The observed distribution is closer to the power-law than to random distribution, corroborating previous analyses (55, 56).

### IV. GRN stability – yeast vs. human

With the off-diagonal *M*_*ij*_’s, the eigenvalues of the matrix *M*_*0*_ can be determined as shown in Fig. 3B for yeast (black dots) and human (red dots) GRNs. Note that the diagonal elements of *M*_*0*_ are set to 0, in comparison with M of Eq. (3). Marked by a vertical line in Fig. 3B, R roughly corresponds to the “radius” of the eigenvalue distribution. The two panels of Fig. 3B also show that the eigenvalue distributions are not noticeably changed by the network structure (random vs. power-law interactions).

The *R* values in yeast and human GRNs differ very slightly (2.2 vs. 1.8 in Fig. 3B) over a wide range of cut-offs used in the estimation. Given that *R* is similar and *D* is 15 times larger in yeast, *R-D* is much more negative in yeast than in human. In other words, yeast GRN is much more stable than human GRN; thus, when perturbed, yeast GRN should be able to return to the equilibrium much more rapidly.

Interestingly, yeast cells can divide 15 times faster than human cells (1.5 hours vs. 24 hours) and, hence, would be perturbed more frequently. It is thus unsurprising that the two GRNs would have different strategies for stability. Yeast GRN may be able to use a simpler strategy for GRN stabilization because unicellular organisms do not have different tissues with different cellular properties. Furthermore, because a typical haploid yeast cell is only 1% as large as an average-sized human cell(57, 58), the transcription rate per unit volume can be much higher in yeast than in human cells. For these reasons, yeast GRN may be able to have non-specific degradation of transcripts that is as rapid as the transcription can keep up. This simple strategy would be neither feasible nor necessary for human GRNs.

In contrast, human GRNs may need to adjust the strategy of stabilization in different cell types. Their larger cell volume also demands far more transcripts; thus, high rate of mRNA degradation may stress the supply to a much larger degree. A suitable strategy for mammals would be to degrade mRNAs more selectively and modestly using miRNAs. Several properties of miRNAs are uniquely suited to this purpose. In a typical mammalian cell, the total number of miRNA molecules is in the same order of magnitude as the number of mRNAs (59, 60). Furthermore, the trunover of miRNA changes much more slowly than mRNAs. The half-life of miRNAs in mammalian cells averages about 120 hours in comparison with that of mRNAs at <10 hours(5, 61). Abundance and slow turnover are the characteristics of miRNAs. In contrast, TFs, the other class of regulatory molecules, have the shortest half lives among mRNAs (see Fig. 3B).

### V. miRNAs and the stability of human GRN

As all repressions, weak or strong, contribute cumulatively to the stability of GRN, the actions of miRNAs are biologically meaningful at the level of the whole GRN. The distribution of miRNAs’ repression effects further supports this view.

#### a. Broad distribution of miRNAs’ degradation effect

The total repression effect is distributed among many miRNAs, most of which are lowly expressed. The repression effect of each miRNA is further distributed over hundreds of target genes, resulting in diffuse repressions over the entire GRN. Here, we evaluate various distributions of miRNA targeting but keeping their aggregate effect constant, at 10% of the total (i.e. 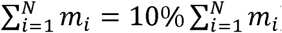,). Fig. 4A shows that, given the same network complexity, the GRN is more likely to be stable when miRNA targeting becomes more diffuse (i.e, targeting more genes with less intensity; see Methods). When only 1% of the mRNAs are targeted for repression by all miRNAs, the probability of GRN stability is only slightly higher than a GRN without any repression. On the other hand, when the repression already covers 25% of all transcripts, further spread would have only incremental benefits to the stability.

**Figure 4.**
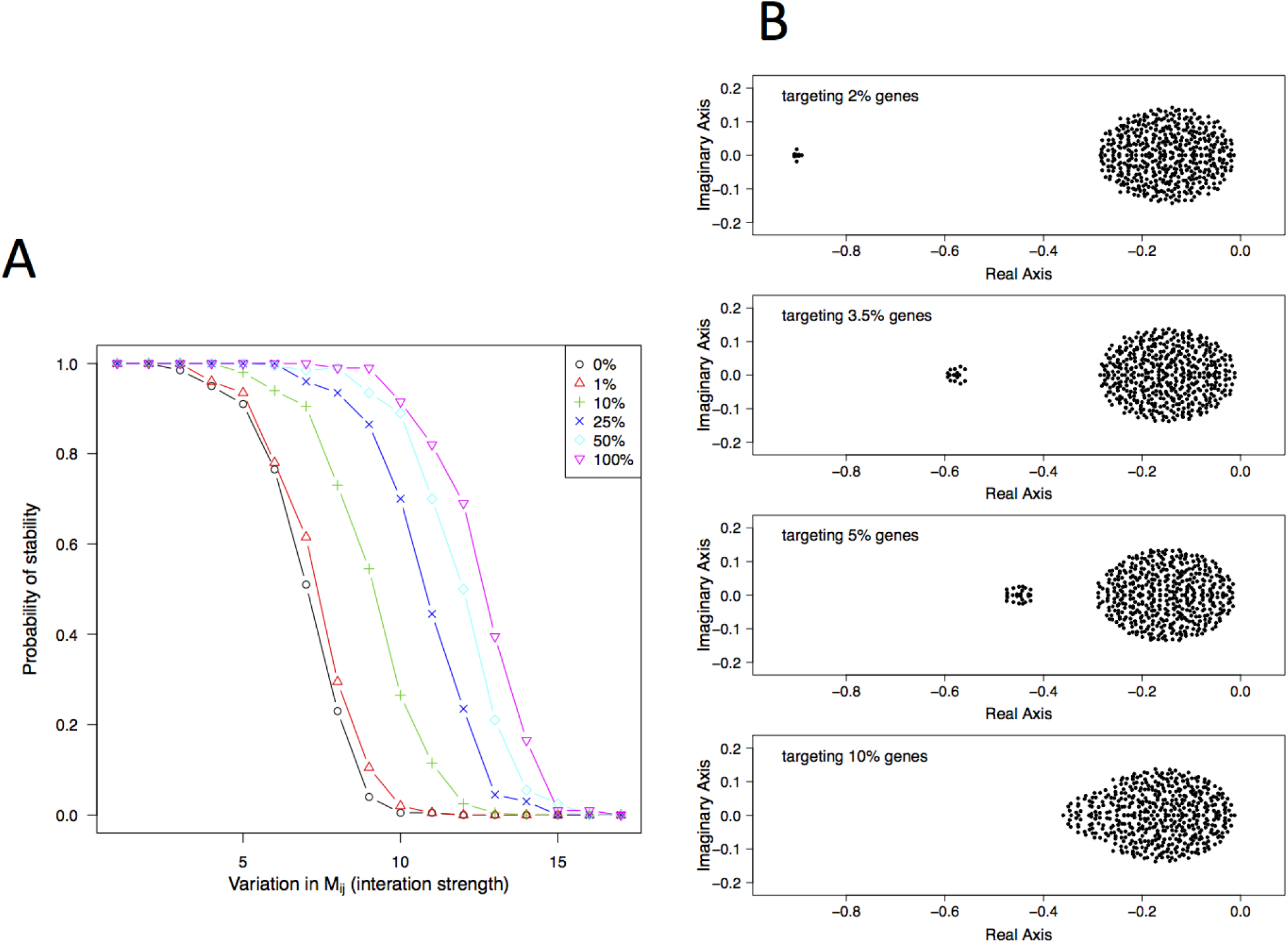
GRN stability in relation to the spread of total miRNA repression in the GRN. (A) The Y axis is the probability of GRN stability, determined by the proportion of cases yielding a negative leading eigenvalue in 200 simulations. The X-axis is the variation in interaction strength presented as the (relative) standard deviation of *M*_*ij*_. The repression is distributed over 1% to 100% of the entire GRN. While the total repression is constant, the probability of stability increases when the effect is spread more broadly over the network. The increase is most rapid from 1% to 25% and slows down gradually. (B) Distributions of eigenvalues as miRNA targeting becomes more diffuse. If the repression is concentrated on a few genes, only a small fraction of eigenvalues is affected, shown by the outliers on the left. Neither the bulk of the distribution nor the leading eigenvalue is noticeably changed. Only when the targeting is sufficiently broad would the entire distribution shift to the left, thus dragging along the leading eigenvalue.

An intuitive explanation is illustrated in Fig. 4b. When miRNAs target a small percentage of genes, only a few eigenvalues are affected and shifted very far to the negative side. The leading eigenvalue is hardly affected, hence resulting in only marginal improvement in GRN stability. The more diffuse the targeting, the more eigenvalues are shifted to the left, eventually dragging the leading value down. Estimates of miRNA targeting fall in the range of 25% −60% of all mRNAs in human cells (43, 44), in reasonable accord with the theoretical prediction of >25%. The next section will explore whether targeting is randomly distributed among all mRNAs.

#### b. Avoidance of very highly expressed mRNAs

If miRNAs function to stabilize GRNs, they are expected to avoid targeting very highly expressed for reason of efficiency. Highly expressed genes can act as “sponges”(62), leaving few miRNAs available for a much larger number of moderately-expressed targets. Furthermore, highly expressed genes are generally less affected by stochastic fluctuations and may suffer less without the stabilizing effect of miRNAs. Fig. 5 is a typical example in which relatively few highly expressed genes have a high number of target sites(Fig. S4).

**Figure 5.**
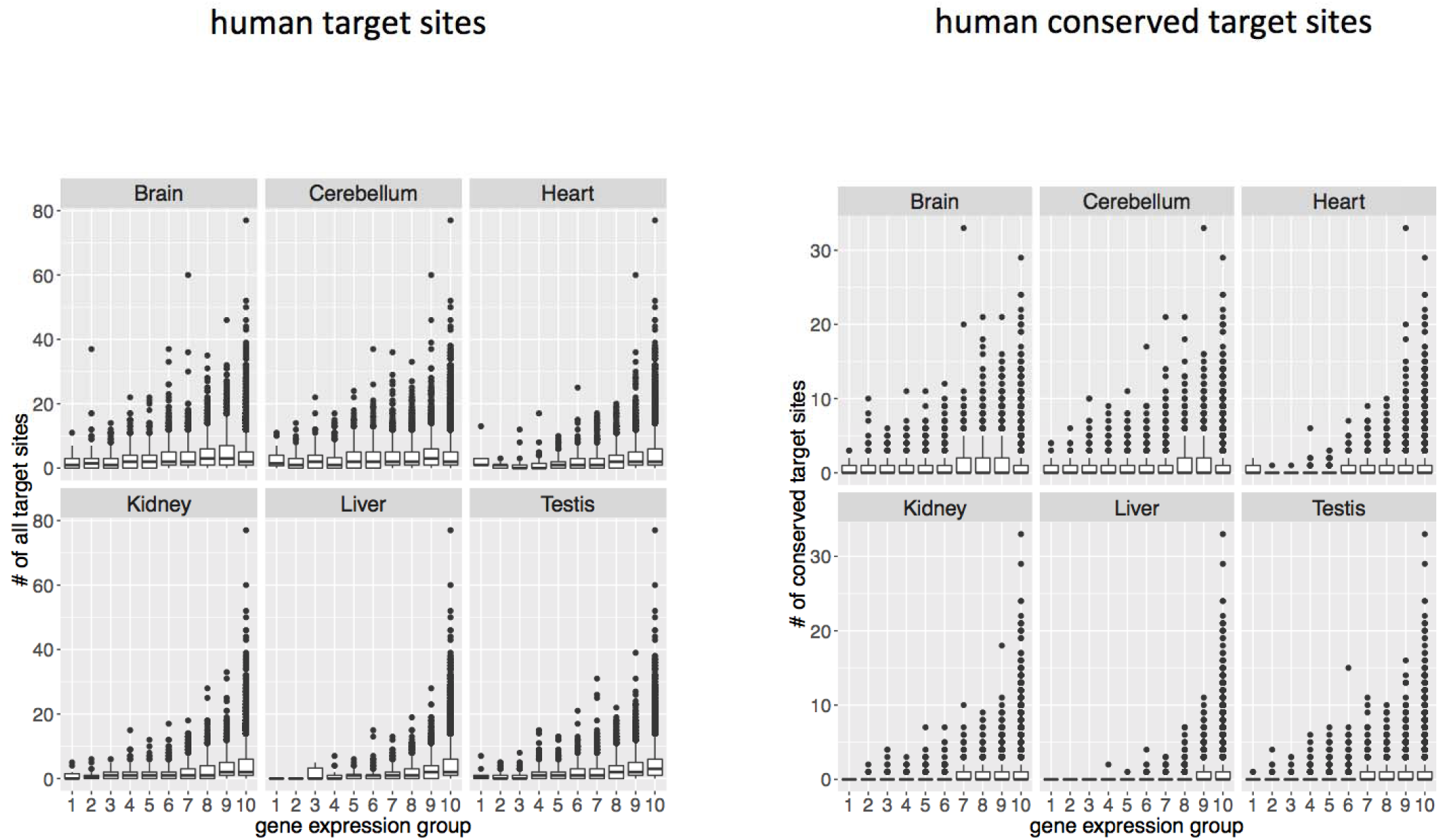
Number of miRNA target sites on genes with different levels of expression, ranging from high to low from left to right in 10 different groups, each containing 10% of all genes. The left set of panel are analysis of all targets of 109 miRNA seeds, the right set of panel are that of conserved targets. Analyses of two different level of evolutionary conservation are shown but the pattern is observable in all (see Supplement). For each level, 6 tissues are analyzed. Note that very highly expressed genes appear to avoid having a very large number of target sites.

#### c. Preference of targeting TFs

GRNs, like many other complex networks, have a hierarchical structure in which nodes of the higher rank regulate those of the lower rank, but not vice versa. Stability of such networks can be decomposed into stability of sub-networks at different hierarchies(63). For GRNs, TFs form a sub-network at a higher hierarchy above non-TFs and are more highly connected(9, 10), which make sub- network of TFs is less stable. Thus, the stability of the GRN would be strongly dependent on the stability of the TF sub-network (Fig. S5-A). It is therefore expected that miRNAs would preferentially target TFs. Indeed, the most conspicuous category of genes significantly enriched for miRNA targets are TFs (64-67). In our re-compilation of miRNA targets in fly, mouse and human, TFs are enriched over the rest of the transcriptome by 16.8%, 15.1%, 13.3%, respectively. If we consider only conserved target genes, the enrichment becomes 89.2%, 49.6% and 41.2% (Fig. S5-B).

## CONCLUSION

The pervasive weak action of miRNAs has been a contentious issue, giving rise to the view that most targets are biologically irrelevant(13, 14). Since the sum of weak repressions accounts for >90% of miRNAs’ total activities it is difficult to reconcile this view with the simple calculation Instead the May-Wigner theory suggests that weak repressions can cumulatively contribute to GRN stability. Furthermore, the more diffuse the repression effect, the more stable the network.

In animals, miRNAs may be the true system-level regulators. It is their wiring pattern, rather than specific links between genes, that is germane to their function. Importantly, by stabilizing GRNs, miRNAs would stabilize the downstream phenotype, albeit indirectly. These molecules are hence the likely agents of developmental canalization proposed by Waddington more than 60 years ago (17-21). An accompanying study (Liufu et al.) found that miRNAs often control the same phenotypes incoherently through multiple target genes. Incoherent control loops are usually associated with stasis, rather than change(28-31). Finally, the contrast between the diffuse actions of miRNAs in animals and the more concentrated repressions in plants (68, 69) raises interesting questions about the divergence between plants and animals in relation to miRNAs’ ancestral functions.

## Author Contributions

Conceptualization, C.W., Y.C. and Y.S.; Methodology, Y.C., C.W., S.A.; Formal Analysis, Y.C.; Writing -Original Draft, Y.C., C.W. and Y.S.; Writing -Review & Editing, C.W., Y.C., Y.S. and S.A.; Supervision, C.W. and Y.C.

## Acknowledgments

We thank Ding Tong, Jacopo Grilli, Si Tang, and Gyuri Barabas for helps and comments in various stages of manuscript preparation. This work was supported by 985 Project (33000-18821105), the National Key Basic Research Program of China (2014CB542006), the Science Foundation of State Key Laboratory of Biocontrol (SKLBC16A37) and The University of Chicago Comprehensive Cancer Center Pilot project.

